# Chemical coupling of biological magnetosomes with amine-functionalized therapeutic antibody conjugates

**DOI:** 10.1101/2025.02.10.637426

**Authors:** S.M Payan, L Gandarias, R Avellan, D Faivre, S Prévéral

**Author notes:** Corresponding author information: Sandra Prévéral : Sandy Marie-Claude Payan Lucia Gandarias Damien Faivre Romain Avellan.

## Abstract

**Background:** Immunotherapy, particularly in cancer treatment, can be enhanced using antibody-drug or antibody-radionuclide conjugates. These conjugates disrupt cell signaling and induce cell death, requiring the targeted antigen to be highly expressed on tumor cells to avoid damage to healthy tissues. A promising strategy to improve delivery is the use of magnetosomes, biological magnetic nanoparticles, which can be guided to tumor sites using magnetic fields. However, most antibody-drug or antibody-radionuclide conjugates are functionalized using the antibody amine groups of the lysine residues on the heavy chains. Therefore, these amine groups are no longer available to bind the antibodies to the magnetosomes.

**Results:** Here, we explore an alternative approach to bind amine-functionalized antibodies to magnetosomes. Using SulfoSuccinimidyl-4-(N-Maleimidomethyl)Cyclohexane-1-Carboxylate (S-SMCC) as a crosslinker, the antibodies are chemically attached to the magnetosome membrane via thiol groups through antibody partial reduction. Our results demonstrate that this chemical process preserves the integrity and functionality of both magnetosomes and antibodies. Moreover, we prove that the produced magnetosome-antibody conjugates are stable under various *in vivo*-like conditions.

**Conclusion:** This coupling method offers significant advantages, enabling the coupling of therapeutic molecules to antibodies combined with the magnetic properties of magnetosomes. This new strategy aims to improve cancer therapy through targeted delivery and rapid accumulation at tumor sites.

## INTRODUCTION

By the end of 2022, 59 monoclonal antibodies have been approved by the FDA for clinical use against cancer. ^1^ Monoclonal antibodies can also be used to deliver drugs or radioactive elements to the tumor site to improve the efficiency of the overall cancer treatment, resulting in a synergy between chemotherapy or radiotherapy and immunotherapy. ^2–4^ These antibody-drug or antibody-radionuclide conjugates have two actions on cancer cells. On the one hand, they disrupt cell signaling, and on the other hand, they induce cell death either through drug cytotoxicity or through radionuclide-induced DNA-damage due to alpha or beta emission. However, the selected antigen for antibody targeting needs to complete several requirements. ^5^ In particular, the targeted antigen needs to be highly expressed on tumor cells with no or very low expression on healthy cells to avoid damage to healthy tissues during both chemotherapy and radiotherapy. ^6^ In addition, to be fully efficient, radionuclide antibody conjugates need to reach the tumor quickly due to the short half-life of therapeutic radionuclides. ^7^ One strategy to overcome kinetic and targeting issues is to improve chemo- or radio-immunotherapy delivery to the tumor through the use of magnetic nanoparticles. ^8,9^ Magnetic nanoparticles have indeed the potential to be magnetically guided to the area of interest using magnetic fields that will improve delivery time to the tumor site. ^10^

In this context, it has been demonstrated that magnetosomes, biological magnetic nanoparticles, have the ability to reach the tumor site 5 minutes after systemic delivery. ^11^ Magnetosomes are magnetic nanoparticles synthesized by magnetotactic bacteria (MTB). ^12^ They are composed of a magnetite crystal covered with a proteolipidic bilayer. Due to a unique set of properties, magnetosomes have been proposed for several biomedical applications. ^13^ In particular, their performance for diagnostic imaging as contrast agents in magnetic resonance imaging ^11,14,15^ and as trace materials in magnetic particle imaging ^16^ surpass that of conventional synthetic particles. Magnetosome have also been proposed as heating agents in magnetic hyperthermia ^17–19^ and photothermia ^19^; and as radio-enhancers in radiotherapy. ^20^ Moreover, the proteolipidic membrane is an inherent advantage of magnetosomes when compared with other chemically synthesized magnetite nanoparticles as it confers stability, prevents aggregation, and more importantly in our case, presents functional groups that can be modified for example with monoclonal antibodies. Magnetosomes have also been investigated as drug delivery carriers as they can be guided using magnetic field gradients towards the target regions where they release their payload. ^21,22^ They also have been tested as mediators for antibody delivery. ^23,24^ However, the combination of magnetosomes and antibody-drug or antibody-radionuclide conjugates has not yet been explored, although this combination potentially offers significant advantages.

In this context, we propose a strategy to use magnetosomes as vehicles for antibodies conjugated with drugs, radionuclides, or labels to potentially improve their delivery to tumor tissues. The need to establish this protocol, arises from the fact that most antibody-conjugates are functionalized using a bifunctional chelator that links to the antibody amine groups of the lysine residues on the heavy chains. ^2,25,26^ Therefore, these amine groups are no longer available to bind the antibodies to the magnetosomes and a different strategy needs to be developed. For example, Trastuzumab, a therapeutic antibody that was initially developed to treat HER2-positive breast cancer, has been conjugated on its amine residues with several radionuclide chelators or drug cross-linkers. ^27,28^ One example is TCMC, a radionuclide chelator, which, when bound to ^212^Pb ^29^, has been used with Trastuzumab to combine radiotherapy with immunotherapy for ovarian and prostate cancer treatment. ^27,30^ For drug-associated immunotherapy, Trastuzumab has been associated to the cytotoxic DM1 agent with SMCC crosslinker (called Trastuzumab-emtansine) to improve breast cancer therapy. ^31^ Therefore, attaching these functionalized antibodies to nanoparticles through amine groups is challenging since the space and residues needed for the drug chelator on the amine residues interfere with subsequent amine-dependent binding.

Here, we develop a new protocol to chemically bind antibody conjugates to the surface of magnetosomes using thiol-dependent antibody binding. For this, we use two different amine-functionalized Trastuzumab antibodies (Trastuzumab-TCMC and Trastuzumab-488) in comparison with a non-conjugated Trastuzumab. The first part of this study focuses on the integrity of the magnetosome after the cellular extraction process. We assess membrane protein content, magnetosome aggregation but also the presence of active amine groups on its surface. We then set up a new experimental protocol consisting in a two-step reaction to efficiently bind amine-functionalized antibody to magnetosomes (Mag-ab conjugate) using the heterobifunctional crosslinker sulfosuccinimidyl-4-(N-maleimidomethyl)cyclohexane-1-carboxylate (SMCC). For this protocol, we set up an optimal antibody reduction step, which is required to bind antibodies to magnetosomes through available thiol groups while preserving antibody integrity. Finally, we assess the efficiency and robustness of the binding to validate our magnetosome-antibody conjugate and we determine its compatible use for future *in cellulo* and *in vivo* assays by exposing magnetosome-antibody conjugates to different *in vivo-* like stress conditions.

## RESULTS AND DISCUSSION

### Design of a chemical strategy to bind amine-functionalized antibodies to magnetosomes

The aim of this study is to set up a protocol to attach antibodies to the membrane of magnetosomes by creating bonds between the thiol groups present in the hinge region of the antibodies with the amine groups of the membrane proteins of the magnetosomes (Figure 1). To achieve this chemical binding, we selected sulfo-SMCC cross-linker to perform oriented binding. Sulfo-SMCC is a heterobifunctional protein crosslinker that possesses sulfhydryl- and amine-reactive properties along with an 11.6 Å spacer arm. In several studies, sulfo-SMCC has been used to attach antibodies to solid surfaces through amine groups of lysine residues present in the fragment crystallizable (Fc) region of the antibodies. ^32–35^ In the present work, we used sulfo-SMCC to bind the amine groups of magnetosome-membrane proteins with the thiol groups of the hinge region of antibodies. To verify the effectiveness of our method, we tested it using three different antibodies: native Trastuzumab antibody (TZ) (with no amine functionalization) as a control, and Trastuzumab-TCMC (TZ-TCMC) and Trastuzumab-488 (TZ-488) as amine-functionalized antibodies (Figure 2).

**Figure 1:**
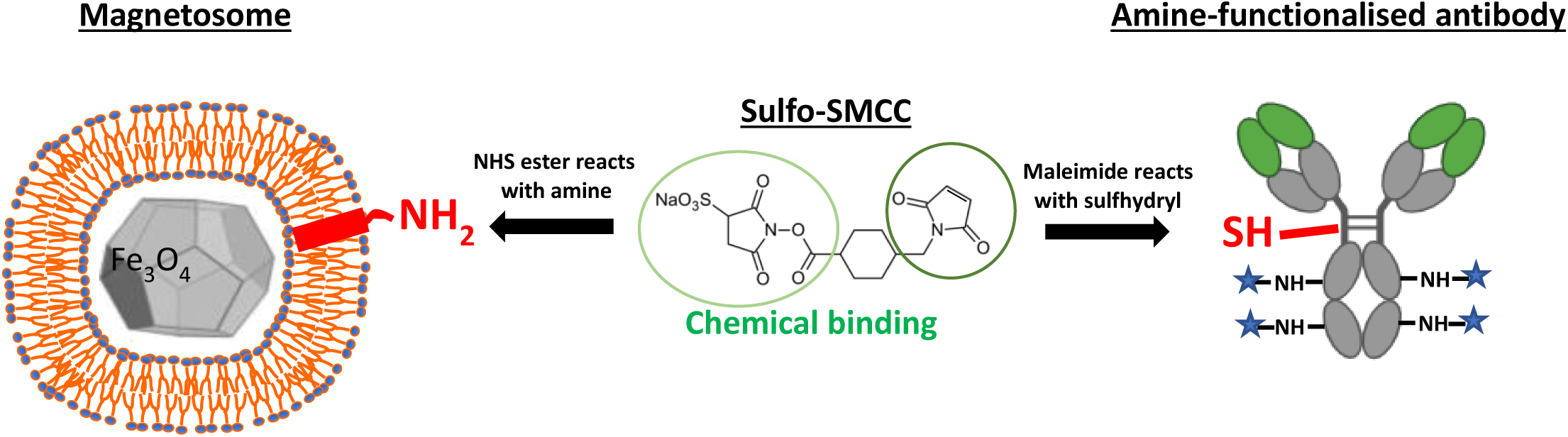
Design of an efficient chemical strategy to bind amine-functionalized antibodies to magnetosomes. Sulfo-SMCC cross-linker was used to bind thiols of partially reduced antibodies with amine groups of magnetosome proteolipid membrane. Created in Biorender.com.

**Figure 2:**
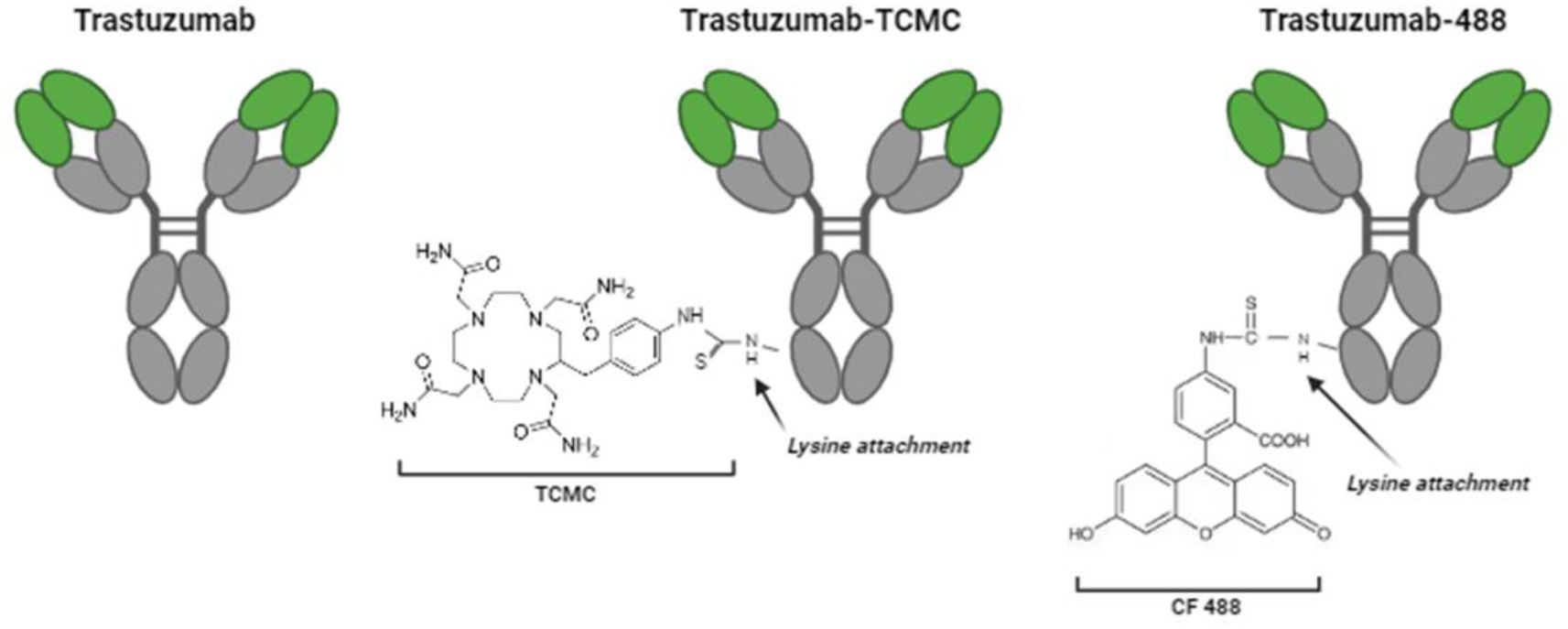
Schematic of the three different trastuzumab antibodies used in this study. Trastuzumab from Proteogenix (left panel) as control antibody, Trastuzumab-TCMC from ORANOmed (middle panel) and Trastuzumab-488 (right panel) as amine-functionalized antibodies. Created in Biorender.com

### Magnetosome characterization

High pressures are used to extract the magnetosomes from the bacteria (see details in the Materials and Methods section). Therefore, before starting to establish the protocol for antibody binding to magnetosomes, we performed a batch characterization and assessed the integrity of the crystal and the membrane of the purified magnetosomes. The magnetic crystal size was determined by TEM imaging, an average size of 57 ± 10 nm was measured (Figure 3A and B) which is consistent with reported crystal size from previous studies. ^11,36,37^

**Figure 3:**
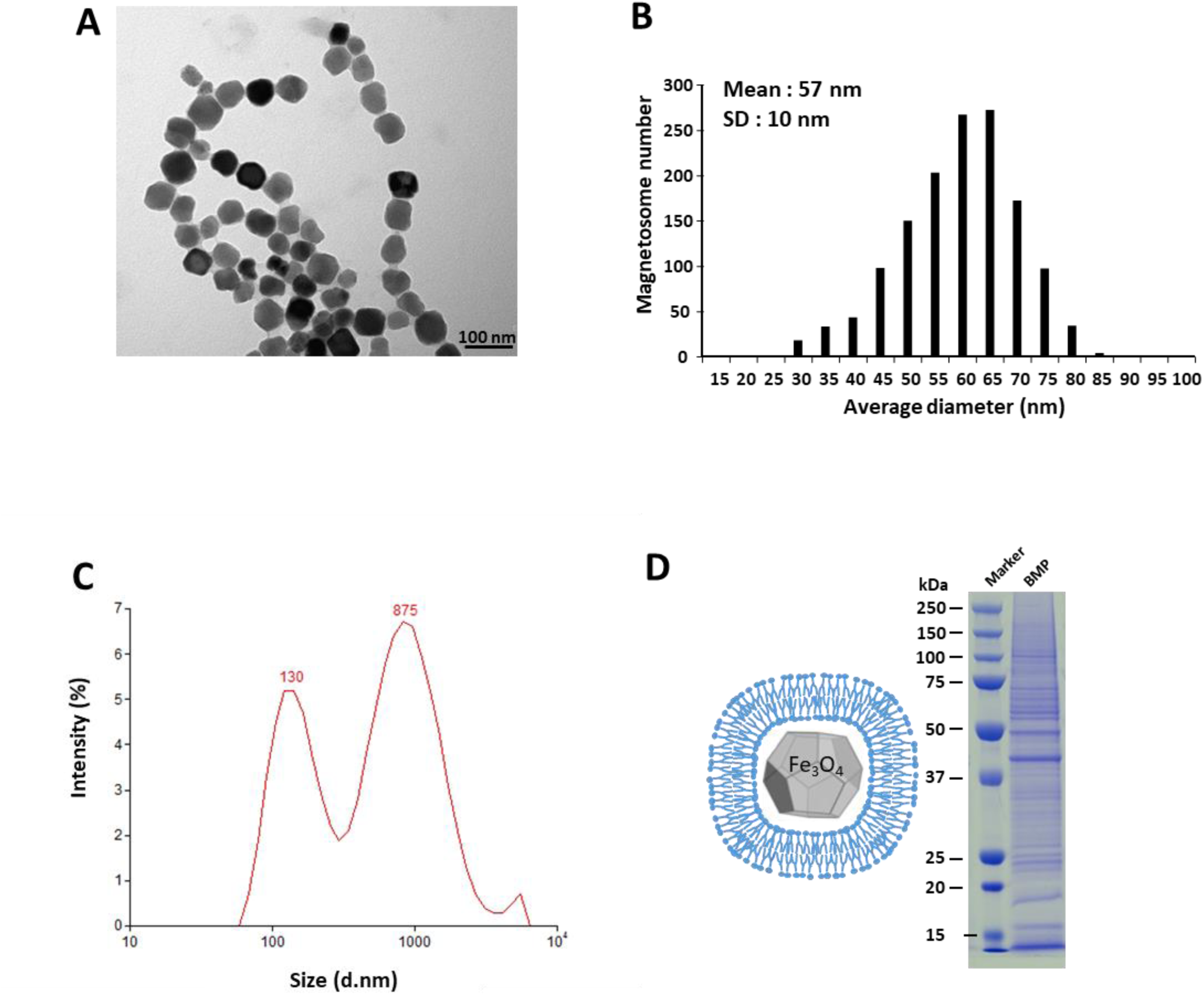
Magnetosome characterization. (A) TEM images of purified magnetosomes. (B) Average diameter (nm) of purified magnetosome from AMB-1 bacteria (n = 1354) determined with Fiji software on TEM images. (C) Magnetosome size distribution as measured with DLS (D) Schematic of a magnetosome (left) and SDS-PAGE analysis of purified magnetosome protein content (right).

The aggregation of the magnetosomes in suspension was determined by dynamic light scattering (DLS). The hydrodynamic diameter profile shows a bimodal distribution with a peak around 130 nm corresponding to isolated or small clusters of magnetosomes and a second peak around 900 nm corresponding to agglomerates of magnetosomes (Figure 3C). It is worth mentioning that the hydrodynamic diameter measured by DLS is the diameter of a sphere that has the same translational diffusion coefficient as the particles in suspension. Importantly, this characteristic is influenced not only by the size of the core of the nanoparticles measured for example by TEM but also by the structure and the properties of the particle surface. The magnetic core of magnetosomes is covered with a proteolipid bilayer that confers them with stability and prevents their aggregation. The presence of this membrane was first confirmed by the zeta potential. Briefly, the zeta potential is an indication of the colloidal stability of a suspension based on the electrostatic repulsive and attractive forces. If the particles in suspension have a large positive or negative zeta potential, they will repel each other and they will not form aggregates. In general, a particle suspension is considered to be stable if it has zeta potential values higher than +30 mV or lower than -30 mV. The zeta potential of the magnetosome suspension was negative and with a mean value of -32 mV, which implies that the electrostatic interactions between magnetosomes are limited.

The integrity of the membrane and the presence of membrane-proteins was further studied by SDS-PAGE (Figure 3D). Compared to the previously described protein profile of AMB-1 magnetosomes, ^38,39^ we demonstrated the conservation of the membrane proteins after the purification process.

As a final control required to carry out the binding protocol, we assessed the presence of primary amine residues in the surface of magnetosomes using a colorimetric detection with the TNBSA reagent. We were able to detect 25.8 ± 8.7 nmol of primary amine residues per µg of iron in the magnetosome suspension.

Together, these results demonstrate that the extraction method used to purify magnetosomes did not affect their crystal and membrane integrity. Furthermore, the amine detection experiments demonstrate the persistence of available amine groups on the magnetosome membrane, essential for their functionalization with antibodies.

### Chemical binding of antibodies to magnetosomes

In order to bind the antibodies to the magnetosome membrane proteins using the SMCC crosslinker, the antibodies need to be reduced so that reactive thiol groups are formed in their hinge region. The antibody reduction was performed using 2-MEA as the reducing agent (Figure 4A). First, we determined the concentration of 2-MEA to perform a partial reduction that preserves the integrity of the antibody while producing thiol groups. Two different 2-MEA reducing reagent concentrations (5 mM and 10 mM) were tested on TZ antibody. To determine if thiol groups were formed after antibody reduction, we performed a colorimetric Ellman dosage (Figure 4B). In addition, antibody integrity was assessed by performing non-reducing SDS-PAGE analysis (Figure 4C). When no reducing agent was added to TZ, as a control, no thiol groups were detected (Figure 4B, left table). In contrast, when TZ was reduced using either 5 mM or 10 mM 2-MEA, we detected a concentration of 3 mM and 14.6 mM thiol groups, respectively. These results demonstrate that both concentrations are compatible for the formation of free thiol groups. However, non-reducing SDS-PAGE analysis demonstrates that only 5 mM 2-MEA reduction preserves antibody integrity as the level of intact antibodies slightly decreases for 5 mM and is almost undetectable for 10 mM compared to non-reduced antibodies (Figure 4C, left panel). To ensure that amine functionalisation on TZ antibody has no impact on antibody reduction, we performed the same experiments on TZ-TCMC antibody (Figure 4B and C, right table). The results obtained for TZ-TCMC antibodies are similar to TZ for both concentrations of reducing reagent (Figure 4 B-C, right panel).

**Figure 4:**
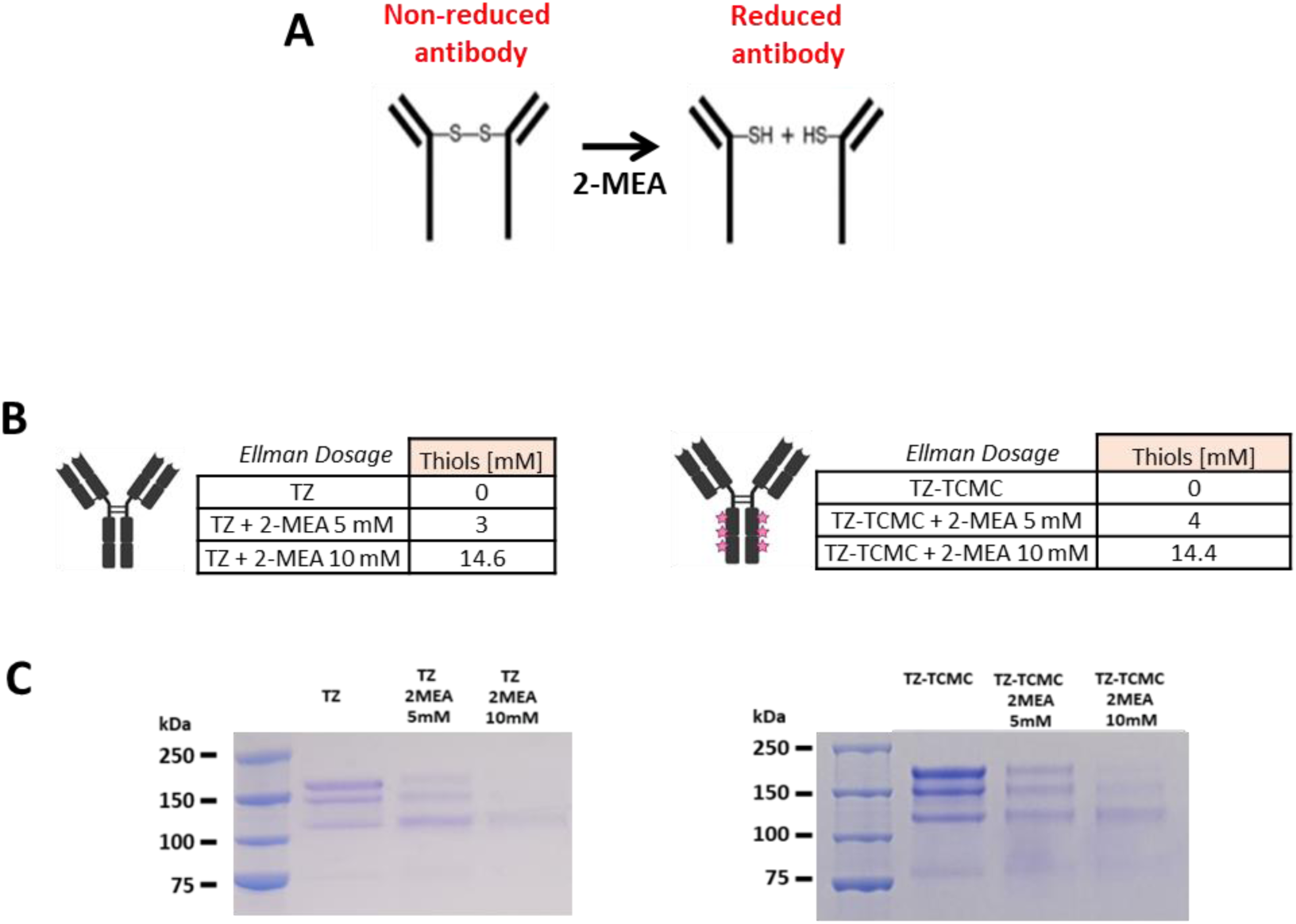
Set-up of antibody reduction protocol. (A) Schematic of antibody reduction using 2-MEA reagent. (B) Ellman dosage analysis of thiols concentration (mM) following partial reduction of Trastuzumab and Trastuzumab-TCMC antibodies at different 2-MEA concentrations (5 mM and 10 mM). Non-reduced antibody was used as positive control. Left panel: Trastuzumab (TZ). Right panel: Trastuzumab TCMC (TZ-TCMC). (C) SDS-PAGE analysis of Trastuzumab and Trastuzumab-TCMC antibody integrity following partial reduction at different 2-MEA concentrations (5 mM and 10 mM). Non-reduced antibody was used as positive control. Left panel: Trastuzumab (TZ). Right panel: Trastuzumab-TCMC (TZ-TCMC).

In view of these results, we decided to use 5 mM of 2-MEA as the reducing agent as, compared to 10 mM 2-MEA, we are able to partially reduce both antibodies forming functional thiol groups while preserving their integrity.

Once the thiol groups are formed in the antibodies they can be bound to the amine groups of the magnetosome-membrane proteins using the crosslinker sulfo-SMCC. As explained on the scheme of Figure 5, the binding reaction of reduced antibodies and magnetosomes is divided into two main stages. The first stage (Figure 5, rectangle 1) consists in the addition of maleimide groups on the magnetosome proteolipid bilayer. These maleimide groups are obtained by the reaction between magnetosomes and sulfo-SMCC, where the NHS ester group of sulfo-SMCC reacts with the amine groups present on the magnetosome surface. To ensure that magnetosome amine groups were functionalized with sulfo-SMCC, we measured the final concentration of free amine groups in Mag-SMCC complex compared to native magnetosome using TNBSA reagent. We detected 15.7 ± 2.5 nmol of primary amine residues per µg of iron in Mag-SMCC. By comparing with the amount detected in Mag (25.8 ± 8.7 nmol per µg of iron), we determine that 39 % of primary amines are functionalized with SMCC in the Mag-SMCC complex. In the second stage of the binding reaction (Figure 5, rectangle 2), the maleimide groups of sulfo-SMCC attached to magnetosomes react with the functional thiol groups of partially reduced antibodies to give the final antibody-functionalized magnetosomes (Mag-Ab) product. Following the protocol established and detailed in the Materials and Methods section, we produced three different types of Mag-Ab conjugates: Magnetosome-SMCC-Trastuzumab (Mag-TZ) as the positive control, and magnetosome-SMCC-Trastuzumab-TCMC (Mag-TZ-TCMC) and magnetosome-SMCC-Trastuzumab-488 (Mag-TZ-488) as magnetosome functionalized with amine-saturated antibodies.

**Figure 5:**
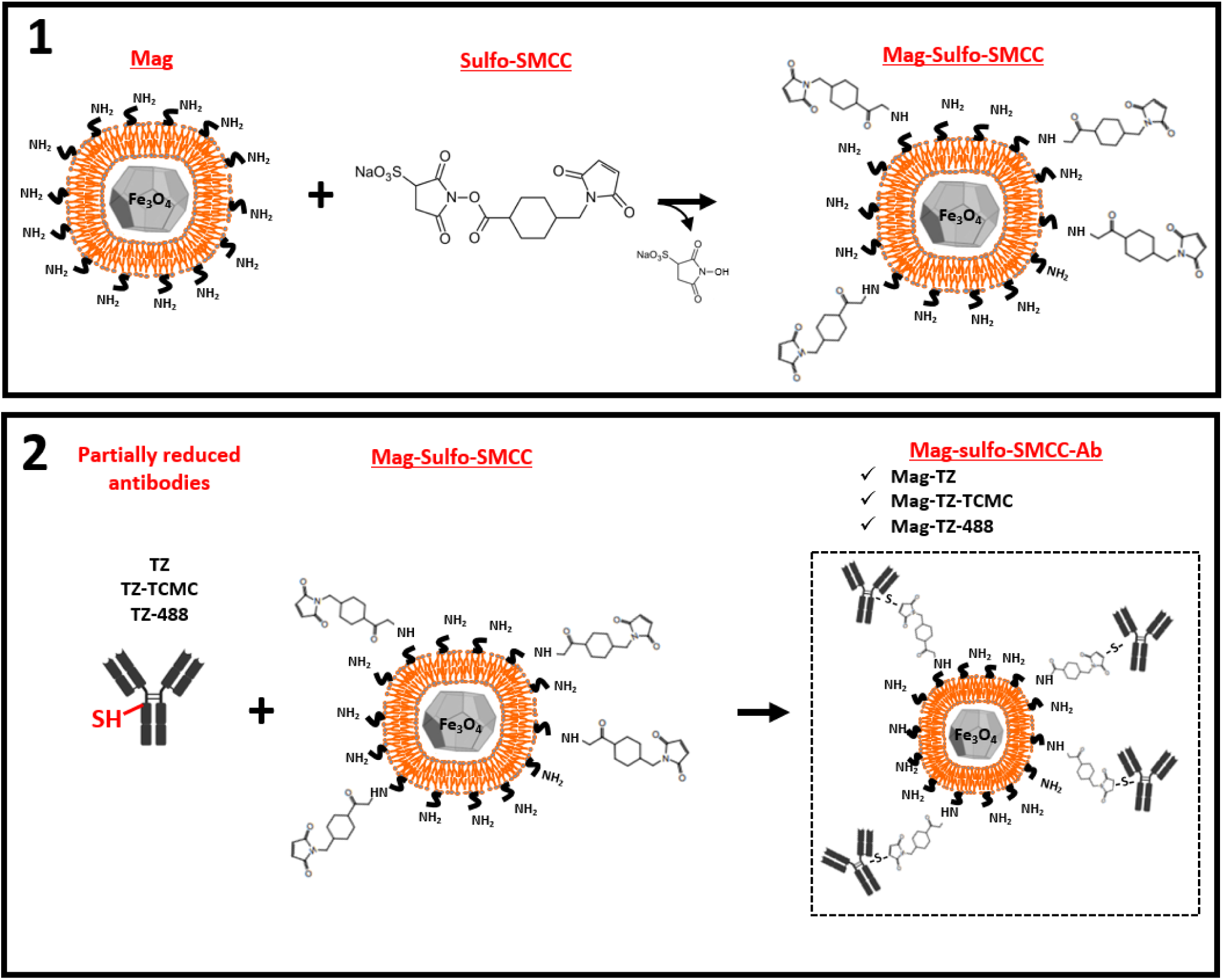
Chemical procedure of antibody binding on magnetosomes (Mag) Schematic of the two-step chemical strategy use for Mag-antibody coupling. The first step (1) is a chemical reaction between a magnetosome (Mag) and a sulfo-SMCC crosslinker that forms maleimide groups on the magnetosome surface (Mag-sulfo-SMCC). The second step (2) is a chemical reaction between partially reduced antibodies (Trastuzumab, TZ; Trastuzumab-TCMC, TZ-TCMC or Trastuzumab-488, TZ-488) and Mag-sulfo-SMCC produced in the first step. This second step leads to the production of our three different magnetosome-sulfo-SMCC-Ab conjugates: Magnetosome-sulfo-SMCC-Trastuzumab (Mag-TZ); Magnetosome-sulfo-SMCC-Trastuzumab-TCMC (Mag-TZ-TCMC) and Magnetosome-sulfo-SMCC-Trastuzumab-488 (Mag-TZ-488).

### *In vitro* Mag-Ab characterization

As a first approach to Mag-Ab characterization, we observed them using TEM (Figure 6A). Mag-Ab show a similar profile when compared to Mag implying that the chemical reactions performed do not affect the structure of the magnetic core of magnetosomes.

**Figure 6:**
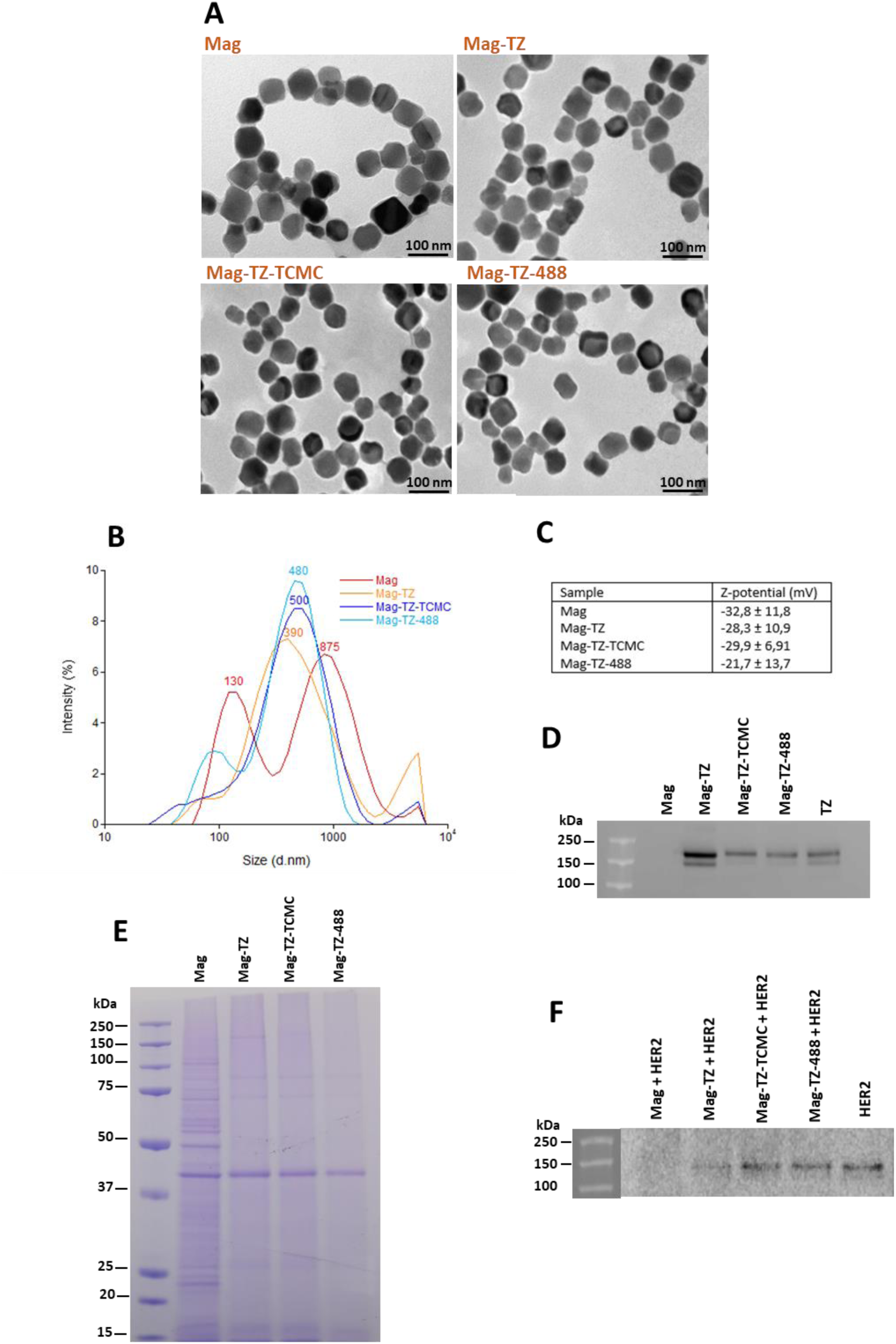
*In vitro* Mag-antibodies conjugates characterization (A) TEM images of purified Mag (upper left) and Mag-antibody conjugates. Mag-TZ (upper right) ; Mag-TZ-TCMC (lower left) and Mag-TZ-488 (lower right) (B) Magnetosome size distribution measured with DLS for control magnetosomes (Mag) compared to Mag-antibody conjugates (Mag-TZ ; Mag-TZ-TCMC ; Mag-TZ-488) (C) Zeta potential values for magnetosomes (Mag) and Mag-antibody conjugates (Mag-TZ ; Mag-TZ-TCMC ; Mag-TZ-488) (D) Western blot detection of Trastuzumab antibody binding in Mag (negative control) compared to Mag-antibody conjugates (Mag-TZ ; Mag-TZ-TCMC ; Mag-TZ-488). Trastuzumab antibody (TZ) was used as positive control. (E) SDS-PAGE analysis of magnetosome protein content of Mag compare to Mag-antibody conjugates (Mag-TZ; Mag-TZ-TCMC; Mag-TZ-488). (F) Western blot detection of HER2 antigen binding in Mag (negative control) compared to Mag-antibody conjugates (Mag-TZ; Mag-TZ-TCMC; Mag-TZ-488). HER2 antigen alone was used as positive control.

To evaluate the success of the antibody-magnetosome binding, we first studied the differences in the hydrodynamic diameter of the samples by DLS (Figure 6B). We observe that the hydrodynamic diameter profile of the Mag-Ab compared to Mag shows wider peaks at around 400 nm (Mag-TZ) and at around 500 nm for Mag-TZ-TCMC and Mag-TZ-488. The zeta potential of the Mag-Ab conjugates (Figure 6C table) also differs from control magnetosomes being lower for the three conjugates. The larger hydrodynamic diameter and zeta potential differences of Mag-Ab compared to the control may be due to the modification of the magnetosome surface. Indeed, functionalization with antibodies generates an increase in antibody protein-protein interactions and magnetosome cluster formation. These results suggest that we had successfully attached antibodies to the surface of magnetosomes.

The confirmation of the antibody-magnetosome binding process was further assessed using western blot analysis on the three types of Mag-Ab (Mag-TZ; Mag-TZ-TCMC; Mag-TZ-488) compared to control magnetosomes (Mag). We were able to specifically detect Trastuzumab antibodies in all the constructs (Figure 6D) demonstrating that the binding of amine-functionalized antibodies on magnetosomes using sulfo-SMCC mediated chemical coupling was successful. The integrity of the magnetosome membrane after chemical coupling was analyzed by SDS-PAGE (Figure 6E). We notice that the magnetosome protein content in Mag-Ab complexes differs from that of control Mag without leading to increased aggregates as observed by TEM (Figure 6A). These results are not surprising since the coupling of antibodies to magnetosomes involves chemical reactions that might disturb some membranous proteins.

We next analyzed the ability of Mag-Ab to bind its specific target, HER2 receptors. To decipher if antibody reduction and binding to magnetosomes do not affect the ability of TZ to bind its target, we performed a HER2 binding assay using a recombinant Human ErbB2/Her2 Fc Chimera Protein. We incubated the three types of Mag-Ab with the HER2 receptor and after performing several washes to remove non-bound HER2, we performed a western blot analysis. We detect HER2 protein specifically bound to Mag-TZ, Mag-TZ-TCMC and Mag-TZ-488 but not in the control with non-functionalized magnetosomes (Mag) (Figure 6F).

Overall, these experiments demonstrate that the protocol we established to produce functional Mag-Ab conjugates is successful as it does not alter magnetosome properties and the obtained Mag-Ab conjugates are able to specifically bind to their target HER2. We thus demonstrate that antibody amine-functionalization does not impede the antibodies to link to magnetosomes nor does it affect their binding ability.

With a perspective of using Mag-Ab conjugates as therapeutic agents, we verified the robustness of antibody binding to magnetosomes under different *in vivo*-like conditions. We subjected Mag-TZ, Mag-TZ-TCMC and Mag-TZ-488 to different biological constraints such as temperature and serum-rich environment and then verified the presence of TZ using western blot analysis (Figure 7). We first investigated if long sonication, required to remove maximal aggregates before injection could affect TZ binding to magnetosomes. For the three constructs, Mag-TZ (Figure 7, upper panel), Mag-TZ-TCMC (Figure 7, middle panel) and Mag-TZ-488 (Figure 7, lower panel), sonication (Mag-Ab + Son) does not affect TZ loading on magnetosomes. We then investigated the impact of a biological warm environment at 37°C (Mag-Ab + 37°C) alone or associated with a sonication process (Mag-Ab + Son + 37°C)(Figure 7). Similarly to the previous tested conditions, TZ is present on the three tested Mag-Ab after these stresses. Finally, we replaced the PBS buffer used for previous stress tests by human serum media and analyzed the impact of serum media incubation at room temperature (Mag-Ab + Serum) compared to human body temperature (Mag-Ab + Serum + 37°C)(Figure 7). In both stress conditions, the TZ antibodies are still linked to Mag-Ab.

**Figure 7:**
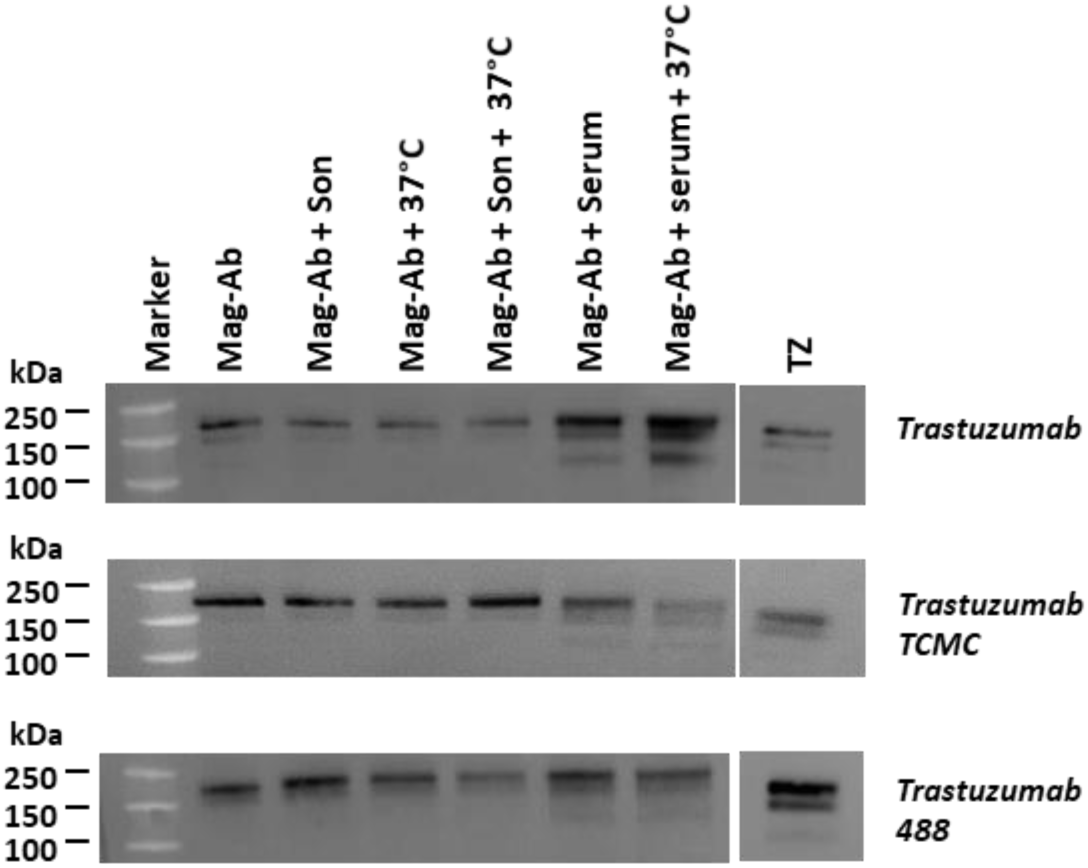
Mag-Ab stability in *in vivo*-like stress conditions Western blot detection of Trastuzumab antibody binding in Mag-antibody conjugates (Mag-TZ : upper panel ; Mag-TZ-TCMC : middle panel ; Mag-TZ-488 : lower panel) under several stress conditions including Sonication (Son), 37°C Heat and Serum. Trastuzumab antibody (TZ) was used as positive control.

In conclusion, we demonstrate that with the established protocol, we can produce functional and robust Mag-Ab conjugates that are not affected by *in vivo*-like conditions and therefore that the constructs are compatible with *in vivo* experiments.

## CONCLUSIONS

In the present study, we demonstrate that amine-functionalized antibodies such as radiolabelled antibodies used for radiotherapy can efficiently be bound to biological magnetosomes through a chemical process. This strategy is based on the use of heterobifunctional crosslinker, Sulfo-SMCC, which binds antibodies to the membrane of magnetosomes using magnetosome functional amine groups and thiol functional groups of partially reduced antibodies. Importantly we demonstrate that the chemical coupling does not affect the properties of either magnetosomes or antibodies.

The combination of antibodies and magnetosomes offers significant advantages to either components taken separately. On the one hand, the magnetosomes become specific as they will bind to the target thanks to the antibodies attached to their surfaces; and on the other hand, the antibodies will more effectively be guided towards the target site by using magnetic field gradients thanks to the magnetic nature of magnetosomes. ^10^ Indeed, according to our previous study, after *in vivo* injection, magnetosomes accumulate in the targeted tumor region in only 5 minutes after injection. ^14^ This is of particular importance for radionuclide-functionalized antibodies that have to arrive to their target quickly due to the short radionuclide half-life. In synergy, Mag-Ab could be used as mediators in adjuvant anticancer-therapies such as immunotherapy, radiotherapy, magnetic hyperthermia and/or photothermia, which will result in a more specific and effective cancer treatment.

## MATERIALS AND METHODS

### Magnetotactic bacteria cultivation and magnetosome purification

The MTB strain *Magnetospirillum magneticum* AMB-1 ^40^ was produced at the “Biomasse-Fermentation” platform of the Marseilles IMM institute. AMB-1 was anaerobically grown in Sartorius BIOSTAT-C30 (30 L) bioreactors until late exponential phase according to the protocol described in ^14^. Bacterial cells were harvested by centrifugation for 20 minutes at 4000 g and stored at −20°C until use. Purified magnetosome suspensions were prepared using the protocol described in Mériaux et al. (2015)^11^. Each 4 g bacterial cell pellets were resuspended in 10 mL of Buffer 1 (20 mM HEPES, 1 mM 2,2’,2’’,2’’’-(Ethane-1,2-diyldinitrilo)tetraacetic acid (EDTA), 8 % glycerol, 0.9 % NaCl, pH 7.5). Bacterial cells were disrupted 3 times with a French press (1200 psi-8.27×10^6^ Pa). Magnetosomes were magnetically purified using a MACSiMAG™ separator (MiltenyiBiotec) and washed 5 times in 10 mL of Buffer 1 and 5 times in 10 mL of Buffer 2 (20 mM HEPES, 8 % glycerol, 0.9 % NaCl, pH 7.5). Magnetosomes were finally resuspended in 200 μL of Buffer 3 (20 mM HEPES, 8 % glycerol, pH 7.5), flash-frozen in liquid nitrogen and stored at −80°C. Magnetosome suspensions were quantified according to their iron content. The iron concentration was determined by Inductively Coupled Plasma Atomic Emission Spectroscopy (ICP-AES) after mineralization in the presence of concentrated nitric acid and overnight incubation at 80°C.

### Antibodies

Trastuzumab biosimilar antibody was purchased from proteogenix (proteogenix, PX-TA1005) and Trastuzumab-TCMC antibody was provided by the OranoMed company. Trastuzumab-488 antibody was homemade produced using Mix-n-Stain 488 Antibody Labeling Kit (Merck, MX488AS50) according to the manufacturer protocol.

### Free amine group dosage

The quantity of free amine groups available on magnetosomes was determined using Picrylsulfonic acid solution 5 % (TNBSA) (Sigma, P2297), which is a colorimetric assay that produces a measurable orange-colored product when it reacts with primary amines. For this, 50 µL native magnetosomes or magnetosomes-SMCC (60 mg.L^-1^) of iron diluted in 0.1 M sodium phosphate buffer (pH 8.5) were distributed in 96-wells plates and mixed with 25 µL 1/500 TNBSA solution previously diluted in 0.1 M sodium phosphate buffer (pH 8.5). After 2 h incubation at 37°C, the reaction was stopped and stabilised by addition of 25 µL SDS 10 % and 12.5 µL of 1N HCl. The absorbance was measured at 335 nm using a 96-well plate reader and the primary amine group quantification was determined using L-alanine amino acid standard and normalized with amount of iron detected by ICP-AES. The mean and Standard deviation was determined (n=6).

### Magnetosome-Antibody (Mag-Ab) chemical coupling

Magnetosomes were functionalized with Trastuzumab, Trastuzumab-TCMC or Trastuzumab-488 antibodies using sulfo-SMCC (Millipore, 573115) cross-linker. The antibodies (12.5 µg) were partially reduced using 5 mM of 2-Mercaptoethylamine hydrochloride (2-MEA, Sigma, M6500) in PBS-EDTA 5 mM at 37°C for 1.5 h. A purification step using Amicon 50K (Millipore, UFC505096) was added to remove the reducing reagent and detached parts of the reduced antibodies. Simultaneously, magnetosomes corresponding to 500 µg of iron were incubated with excess sulfo-SMCC solution (1.36 mg sulfo-SMCC at 5 mg mL^-1^) in PBS for 1.5 h to form maleimide groups using NH_2_ functions on their surfaces. Then, the obtained magnetosomes-maleimides were magnetically washed using the MACSiMAG™ separatorin PBS to remove the excess of sulfo-SMCC solution. Finally, magnetosomes-maleimides were incubated with the partially reduced antibodies for 3 h in 0.1 M Sodium Bicarbonate buffer pH 7 at room temperature. Magnetosomes-antibody conjugates were then magnetically washed 3 times with Buffer 3 and finally resuspended in 100 µL Buffer 3. The final iron concentration was assessed using ICP-AES analysis.

### Free thiols group dosage

The quantity of free thiol groups available in reduced antibodies was determined using Ellman’s Reagent (2-nitrobenzoic acid, Thermofisher #22582), which is a colorimetric assay that produces a measurable yellow-colored product when it reacts with sulfhydryl groups. After 2-MEA reduction and purification step, antibodies were diluted in 0.1 M sodium phosphate pH 8, distributed in 96-wells plates and mixed with 1 mM EDTA reaction buffer and Ellman’s reagent. After 15 min incubation at room temperature, the absorbance was measured at 412 nm using a 96-wells plate reader and the concentration of sulfhydryl groups was determined using a cysteine standard.

### Transmission electron microscopy

Transmission electron microscopy (TEM) was performed on magnetosomes and bacteria deposited onto 200-mesh formvar carbon-coated copper grids and washed three times with MilliQ water. The images were collected using a Tecnai G2 BioTWIN (FEI Company) electron microscope equipped with a charged-coupled device (CCD) camera (Megaview III, Olympus Soft imaging Solutions GmbH) using an accelerating voltage of 100 kV. For crystal size measurements, TEM images were analysed using the Fiji software. The mean and Standard deviation was determined.

### DLS and Zeta potential measurement

Dynamic light scattering (DLS) and zeta potential analyses were carried out on magnetosomes suspended in milliQ water at a concentration of 0.025 µg Fe. mL^-1^. The samples were analysed using a Zetasizer Nano-ZS (Malvern Panalytical) in a disposable sizing cuvette with a backscattering angle of 173 ° at 25°C. The displayed results are the mean values of 12 measurements.

### SDS-PAGE and western blot

The amount of magnetosomes corresponding to 10 µg (for western blot analysis) or 20 µg of iron for total protein analysis) was suspended in 1X LDS sample buffer (Thermofisher, NP0007), boiled for 5 min at 95 °C and loaded into a 10 % Bis-Tris gel (Invitrogen, NP0315BOX). SDS-PAGE was performed in 1X MOPS SDS running buffer (Invitrogen, NP0001). For total protein analysis, the gel was washed twice in distilled water and incubated with Imperial protein stain (Thermofisher, 24615) according to the manufacturer protocol. For western blot analysis, proteins were then transferred in 1X MOPS 20 % ethanol 0.05 % SDS buffer onto a nitrocellulose membrane using a wet transfer system for 1.5 h at 100 V. After a step of saturation (1 h at room temperature in 5 % non-fat powdered milk dissolved in 1X Tris-buffered saline-Tween 0.2 %,(TBS-T-milk)), the membrane was then immunoblotted with primary anti-Trastuzumab (Biotechne MAB95471-SP) or anti-HER2 antibody (Sigma SAB4500789,) diluted by 1000 in TBS-T-milk and a secondary anti-rabbit IgG-Peroxidase (Sigma, A6154) antibody diluted by 10000 in TBS-T-milk. The chemiluminescence signal was obtained with the SuperSignal™ West Pico Chemiluminescent Substrate (Millipore, WBKLS0100) and imaged with a G:BOX imaging system (Syngene).

### HER2 binding analysis

To test the binding ability of Mag-Ab conjugates with their target receptor (HER2), magnetosome suspensions containing 40 µg of iron were diluted in 30 µL phosphate buffer saline (PBS) and incubated with 0.6 µg recombinant Human ErbB2/Her2 Fc Chimera Protein (Biotechne, 1129-ER-50) for 1 h at 37°C. The samples were magnetically washed 3 times in PBS solution to remove non-attached HER2 debris, suspended in 22.5 µL water and analysed using western blot as detailed in the previous section.

### a. Magnetosome-Trastuzumab antibodies stability analysis

Mag and Mag-Ab conjugates corresponding to 15 µg of iron were resuspended in 20 µL PBS or 20 µL of human serum (Sigma, H4522) and sonicated for 10 minutes and/or incubated at 37 °C for 1 h. Then, they were magnetically washed 3 times in PBS and resuspended in 22.5 µL water. Qualitative western blot analysis was performed to assess the presence of Trastuzumab antibodies attached to magnetosomes.

## Aknowledgements

We would like to acknowledge Orano Med company to have provided us with the antibody and Orano for the financial support. L.G. would like to acknowledge the financial support provided through a postdoctoral fellowship from the Basque Government (POS_2022_1_0017).

## Abbreviations

2-MEA: 2-Mercaptoethylamine hydrochloride
AMB-1: Magnetospirillum magneticum
CCD: Charged-coupled device
DLS: dynamic light scattering
EDTA: Ethylenediaminetetraacetic acid
Fc region: fragment crystallizable region.
HER2: human epidermal growth factor receptor 2
ICP-AES: Inductively Coupled Plasma Atomic Emission Spectroscopy
IgG: Immunoglobulin G
LDS: Lithium dodecyl sulfate
Mag: magnetosome
Mag-Ab: magnetosome-antibody conjugates
Mag-SMCC: magnetosomes-Sulfosuccinimidyl 4-(N-Maleimidomethyl)Cyclohexane-1-Carboxylate
Mag-TZ: magnetosome-Trastuzumab
Mag-TZ-488: magnetosome-Trastuzumab-488
Mag-TZ-TCMC: magnetosome-Trastuzumab-2-(4-isothiocyanatobenzyl)-1,4,7,10-tetraaza-1,4,7,10-tetra-(2-carbamoylmethyl)cyclododecane
MOPS: 3-(N-morpholino)propanesulfonic acid
MTB: magnetotactic bacteria
PBS: Phosphate buffered saline
SDS: Sodium dodecyl-sulfate
SDS-PAGE: Sodium dodecyl-sulfate polyacrylamide gel electrophoresis
SMCC: Sulfosuccinimidyl 4-(N-Maleimidomethyl)Cyclohexane-1-Carboxylate
TBS: Tris-buffered saline
TBS-T: Tris-buffered saline-Tween
TCMC: 2-(4-isothiocyanatobenzyl)-1,4,7,10-tetraaza-1,4,7,10-tetra-(2-carbamoylmethyl)cyclododecane
TEM: Transmission electron microscopy
TNBSA: 2,4,6-Trinitrobenzenesulfonic acid solution, picrylsulfonic acid solution
TZ: trastuzumab antibody
TZ-TCMC: Trastuzumab-2-(4-isothiocyanatobenzyl)-1,4,7,10-tetraaza-1,4,7,10-tetra-(2-carbamoylmethyl)cyclododecane antibody
TZ-488: Trastuzumab-CF488 dye

## TABLE OF CONTENT GRAPHICS

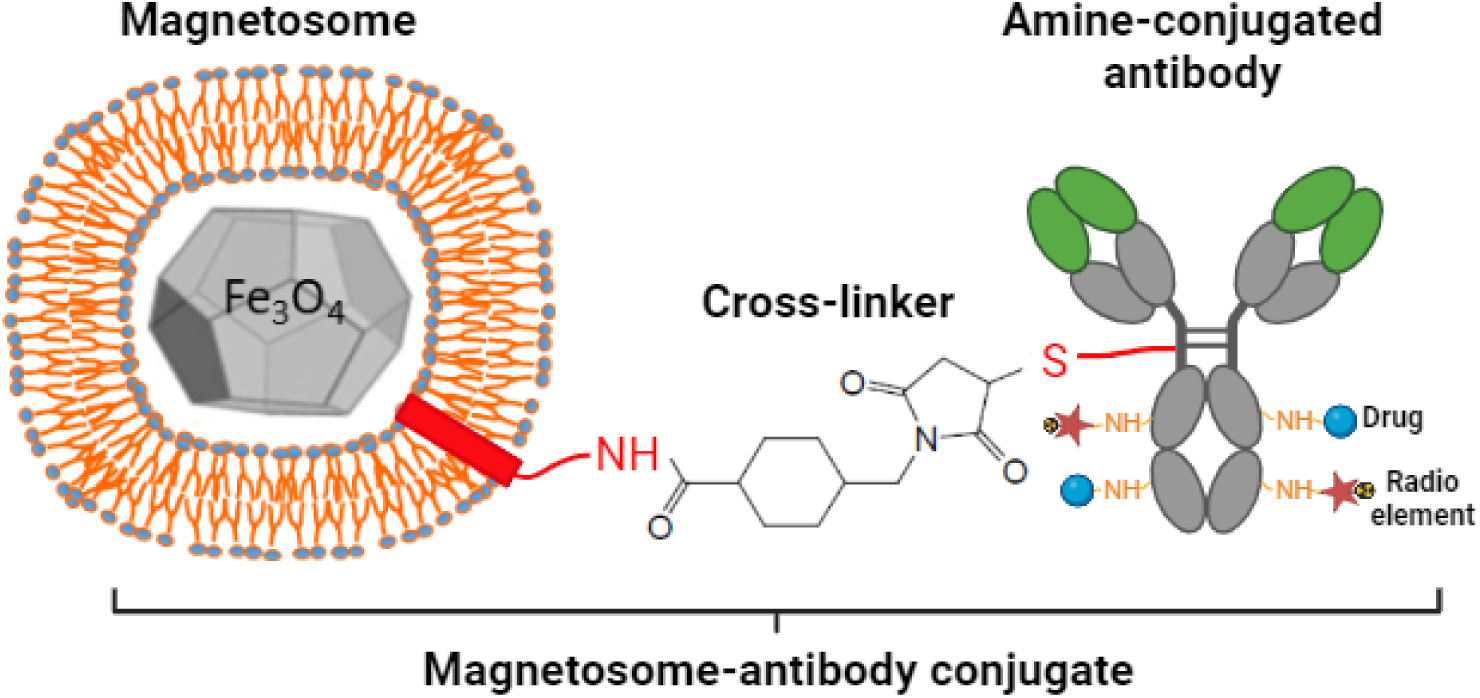

